# Inhibition of GEF-H1-RhoA signaling in inflammation with a stapled peptide mimicry of the RhoA^67-78^ helix

**DOI:** 10.1101/2024.11.18.624118

**Authors:** Clara Gathmann, Tiansheng Liu, Shuhuai Yang, Weitong Cynthia Wang, Edith Chan, Karl Matter, Maria S. Balda, A. W. David L. Selwood

**Author notes:** Authors contributed equally (Clara Gathmann, Tiansheng Liu).

## Abstract

Guanine exchange factors (GEFs) are considered hard to drug with conventional small molecules, they lack conventional deep binding pockets and binding ligands are seldom reported. Here we report the design of a stapled peptide stP5 targeting the interaction between cytoskeletal regulator RhoA GTPase and its activator guanine exchange factor H1 (GEF-H1). StP5 is a modified RhoA mimic based on a previously identified bioactive α-helical epitope to GEF-H1. StP5 effectively inhibits GEF-H1-induced morphological and transcriptional changes in cellular models for inflammation and does not affect the related GEF p114RhoGEF (ARHGEF18). StP5 peptide is approximately 100 fold more active in cellular assays than the unstapled P5 peptide. We provide a bioinformatic analysis of the stP5 bindings site in different GEFs, providing a basis for this selectivity. The GEF-H1 inhibitor stP5 represents a step towards fully drugging GEF-H1.

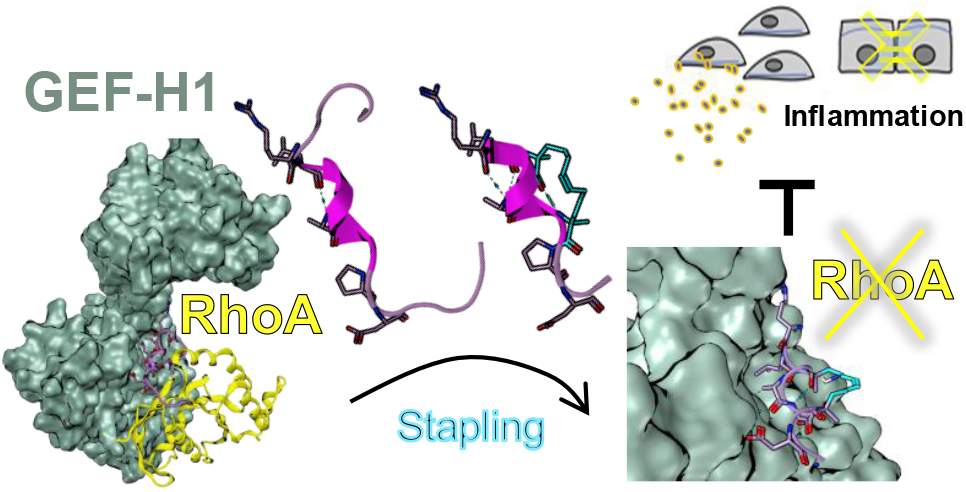

## INTRODUCTION

The Ras family of small GTPases are guanine nucleotide-binding monomeric proteins broadly involved in signal transduction, comprising Ras, Rho, Arf, Rab, and Ran sub-families. They cycle between an inactive GDP-bound and an active GTP-bound conformation, the latter binds and activates effector proteins to relay cell signaling events^[1],[2]^ (SI, Fig. S1). For their role in regulating key cellular processes, Ras GTPases like K-Ras are broadly investigated as therapeutic targets^[3]^. Inhibitors of signaling by other GTPases of the Ras superfamily like Rho are, however, still poorly represented. Rho GTPases act as central cytoskeleton regulators^[4-5]^ and their dysregulation has been associated with a broad range of diseases, including cancer^[6-7]^, fibrosis^[8-9]^, and inflammatory disorders^[10-12]^, making them an attractive signaling mechanism to target.

Direct targeting of the Rho GTPase nucleotide binding site is not viable due to its picomolar affinity to GDP and GTP. A few inhibitors binding allosteric pockets on the Rho subfamily members RhoA, Rac1, and Cdc42 have been reported^[13-17]^, but all remained early-stage candidates. The most advanced clinical strategies targeting Rho thus far relied on indirect targeting their membrane localisation^[18-20]^ or the use of biologics^[21-22]^. There is thus an interest in tackling Rho GTPase signaling through other approaches, including by targeting their interaction with regulators^[23-25]^. The Ras superfamily of GTPases is tightly regulated by guanine exchange factors (GEFs) and guanine activating proteins (GAPs), which accelerate the exchange of GDP for GTP or accelerate the hydrolysis of GTP to GDP, respectively^[26]^ (SI, Fig. S1B). Interestingly, and unlike Ras, Rho GTPase levels rarely alter in disease. Instead, their regulators are often responsible for their over-activation^[23-24]^. Rho GTPases are also strikingly outnumbered by their regulators, with >80 Rho GEFs described for 22 Rho proteins, allowing for tight spatiotemporal regulation of Rho signaling^[27]^. Drugging Rho GEFs may thus offer a targeted therapeutic intervention to tackle Rho signaling. Specifically, we focus on GEF-H1, a GEF reported to over-activate RhoA in disease^[28]^. Notably, GEF-H1 overexpression and increased RhoA activation promotes inflammation and fibrosis in diseases affecting the epithelial and endothelial barrier^[29-31]^, and GEF-H1 overexpression, leading to increased cell proliferation, has been linked to cancer progression in melanoma^[32]^, sarcoma^[33]^ and carcinoma^[34-35]^. Targeting Rho GEFs has little precedent thus far (reviewed in [^[23]^]), perhaps due to the inherent challenges associated with their structures, lacking deep binding pockets, typical for protein-protein interactions (PPIs)^[36]^. Small molecule binders of the Rho GEFs LARG, Akap13, and TrioN have been discovered following docking studies, but all share a PAIN (Pan-Assay InterfereNce) motif, thus requiring further validation^[23]^. Here, we propose an alternative ligand-based design approach to target GEF-H1 by generating a RhoA-derived peptidomimetic. It is known that PPIs can be narrowed down to hot-spot residues that contribute to most of their binding affinity^[37]^. ‘Hot segments’ can be identified by excising epitopes of the natural protein partner and further used for downstream drug design^[38]^. The Zheng group has previously identified the linear peptide region 43-58^RhoA^ in Rho GTPases as a potential hot segment which inhibited the Rho GTPase/GEF interactions in the micromolar range^[39]^. Similarly, our group has previously identified a micromolar bioactive α-helical peptide P5 from region 62-80^RhoA^ based on a screen of RhoA segments in a GEF-H1 overexpression model^[1]^. A membrane permeable version of P5 (TAT-P5) was effective in fibrotic, proliferative, and inflammatory models with RhoA/GEF-H1 activation by disease mediators such as TGFβ1, LPS, or TNFα^[1]^. These studies demonstrate the feasibility of GEF targeting through a RhoA-derived peptide, but still involve long, non-drug-like sequences of >15 amino acids. There is thus interest in further narrowing down these segments to fewer hot-spots and consolidate starting points for future molecular designs.

For epitopes with an α-helical secondary structure, peptide stapling is an attractive method to reinforce helicity and potentially improve physicochemical properties of peptide segments^[40-41]^. P5 was designed from a model of the GEF-H1/RhoA interaction and is derived from the 62-80^RhoA^ α-helix^[1]^ binding GEFs near the frontier between their Dbl homology (DH) and Pleckstrin homology (PH) domains, the main domains responsible for the nucleotide exchange on RhoGTPases (Fig. 1A). An L72A mutation was introduced to reduce steric interactions with GEF-H1-specific H345 residue^[42]^. The central region of the P5 peptide is an α-helix interacting with GEF-H1 through residues R68, L69, A72, and D76 on one face of the helix and with S73 on the other face (Fig. 1A). In our previous study, truncating the peptide down to the 68-77 region resulted in an inactive compound, which we hypothesized was likely due to a loss of secondary structure. In this study we report the improved activity of a constrained “stapled” version of P5 peptide and describe its improved activity in cell-based assays of GEF-H1 function.

**Figure 1.**
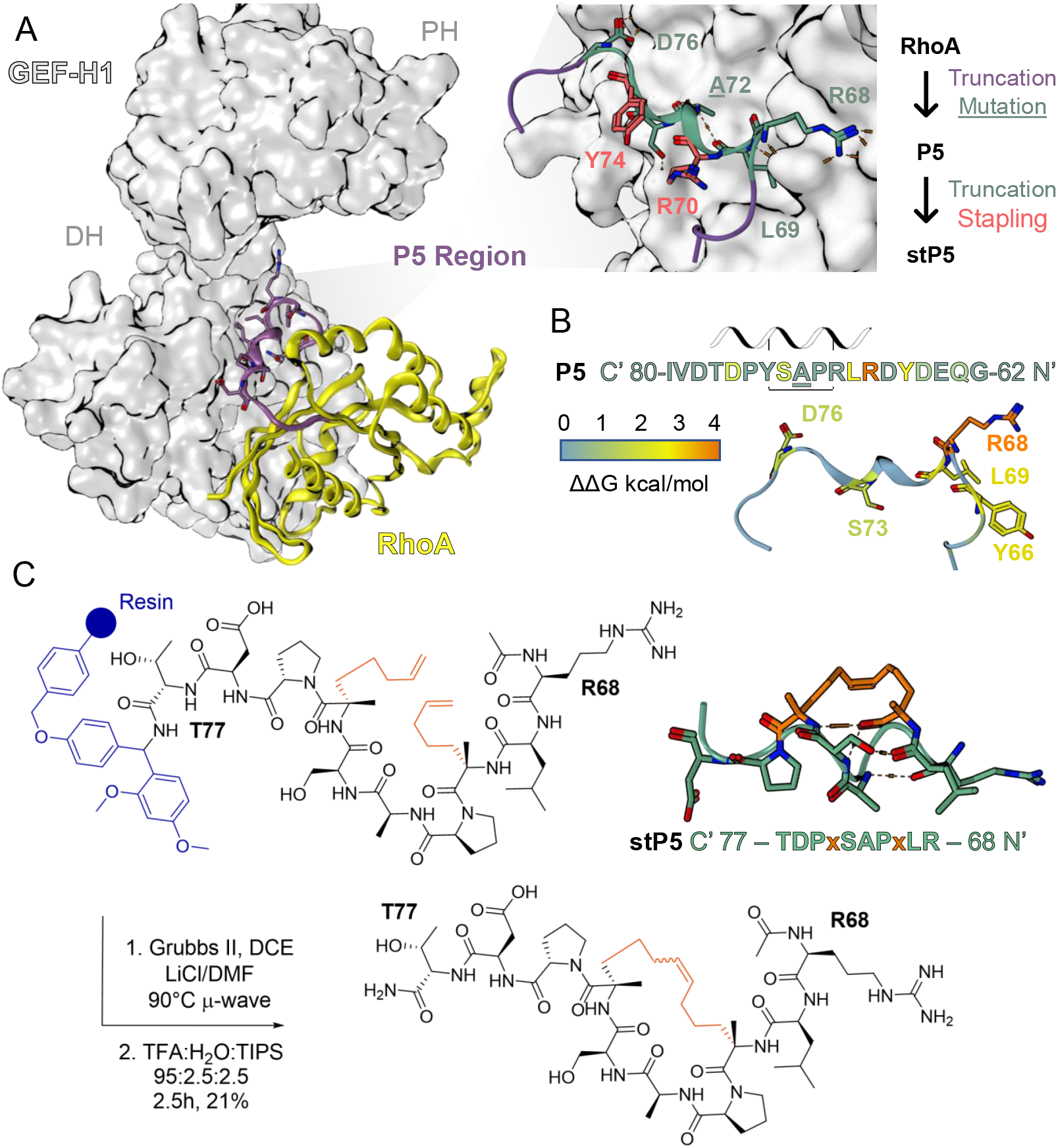
Design and synthesis of a stapled peptide derived from a RhoA epitope. **(A**) A model of the interface between GEF-H1 and RhoA was built based on the Akap13-RhoA crystal structure (PDB 4D0N) by mutation and energy minimisation. The region for the previously designed bioactive peptide P5 is coloured in purple. Right is an enhanced view of peptide P5 in the GEF-H1 model, showing its main residues at the interface to GEF-H1. The helical part of P5 is shown in green and suggests truncation of the purple ends. The two solvent-exposed residues R70 and Y74 suggest suitable stapling points. An L72 to A72 mutation was introduced due to potential clashes with the GEF-H1 structure. (**B**) Alanine scanning mutagenesis results of peptide P5 in complex with a GEF-H1 model. Binding hot-spots are coloured orange and yellow, whereas parts of the peptides unaffected are coloured in blue. The full results are presented in the supporting information section B. (**C**) Resulting design and synthesis of stapled peptide stP5. An 8-carbon long staple was used and introduced by ring-closing metathesis (RCM) on the commercial, on-resin, ring-open peptide, followed by acidic cleavage from the resin.

## RESULTS AND DISCUSSION

Inspection of the binding mode of P5 in a GEF-H1 homology model suggests solvent-exposed Y74 and R70 might be appropriate stapling points to enforce the helicity of the 68-77^RhoA^ fragment (Fig. 1A right). Alanine scanning mutagenesis of the P5 region in the GEF-H1/RhoA model using the Robetta software^[43]^ further confirmed the suitability of these residues as binding energy was not impacted by their mutation (Fig. 1B). Robetta predicts that the biggest binding hot spots are mostly located within the helical region of P5, with R68 producing a remarkable shift of 4.2 kcal/mol (extended data in SI, Table S2). An *i*-*i+4* stapling method using the well-established (S)-pentenylalanine hydrocarbon linkers^[44]^ was thus chosen for stapling a truncated form of P5, resulting in an 8-carbon-long staple. StP5 was synthesized from its ring-open form immobilized on a Rink amide resin, using Hoveyda-Grubbs 2^nd^ generation catalyst, microwave-assisted ring closing metathesis adapted from the method described by Jackson *et al* ^[45]^ (Fig. 1C).

We first examined the helical content of the stP5 using circular dichroism (CD) and compared it to the unstapled peptide as control (SI, Fig. S2). Both unstapled and stapled peptides showed a low proportion of helical content (8.2 and 9.9% respectively) in 50% trifluoroethanol^[46]^. However, when we examined the ability of stP5 to bind GEF-H1 *in vitro*, improved binding to GEF-H1-DHPH was observed. Steady-state analysis surface plasmon resonance (SPR) against the GEF-H1-DHPH fusion protein immobilized on a CM5 chip shows that peptide P5 binds GEF-H1 with an affinity (K_D_) of 100 μM (Fig. 2A), whereas stP5 has a K_D_ of 34 μM (Fig. 2B). Thus, direct binding of both peptides to GEF-H1 was demonstrated and stapling improved the affinity of the P5 against GEF-H1 3-fold while shortening its sequence considerably.

**Figure 2.**
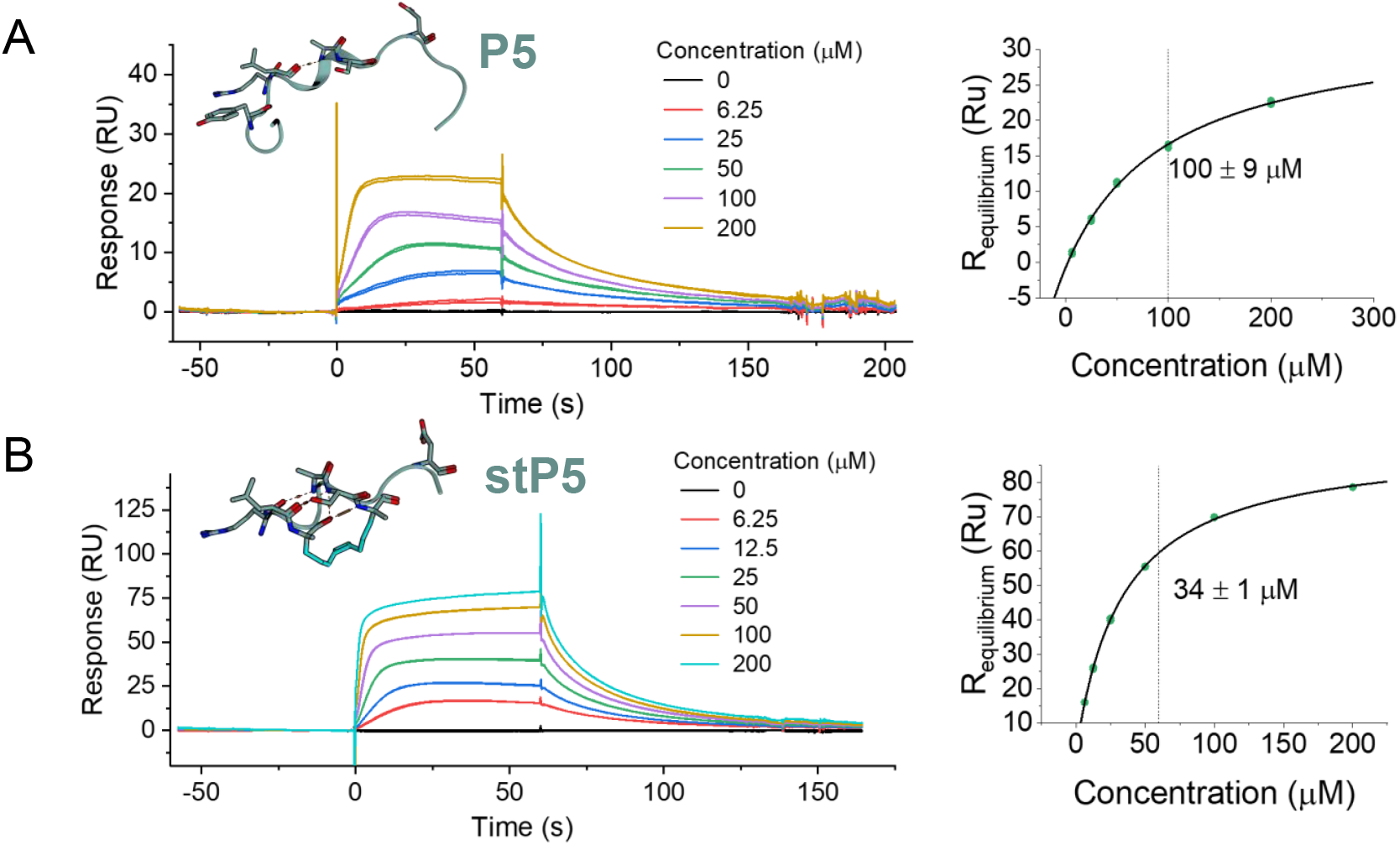
Affinity of P5 and stP5 by surface plasmon resonance (SPR) against the DHPH domains of GEF-H1. GEF-H1-DHPH was immobilized by amide coupling on a CM5 chip. Different concentrations were flown over the surface of (**A**) P5 (4200 RU chip) and (**B**) stP5 (8000 RU chip) and their binding affinities (K_D_) were determined by a non-linear fit of equilibrium responses against concentration. Experiments were run in duplicates with K_D_ ± SD of the concatenated mathematical fit.

We next investigated if the stP5 Rho mimic was active in cellular models of GEF-H1 overexpression. StP5 was tested in an MDCK cell line stably expressing VSV-tagged GEF-H1 regulated by a doxycycline-inducible promoter, as described previously^[1]^. Activation of RhoA by GEF-H1 results in pronounced cytoskeletal rearrangements, marked by stress fiber formation and transcriptional activation^[28]^. As a consequence of the cytoskeletal changes, cells become more elongated and variable in size, which can be assessed by quantification of the cell aspect ratio and its cell-to-cell variability. We compared the effect of TAT-P5, our previously described GEF-H1 inhibitor^[1]^, to stP5 normalized to untreated cells. We also investigated stP5 conjugated to the TAT element (obtained commercially) to improve its cellular penetrability. MDCK cells treated with doxycycline and TAT-P5 (10 μM), stP5 (1 μM) or TAT-stP5 (1 μM) showed improved junction formation and reduction of the elongated phenotype visible upon GEF-H1 induction (Fig. 3A). A 0.1 μM treatment of stP5 reduced cell shape changes by about 60%, comparable to 80% reduction with a 10 μM TAT-P5 treatment (Fig. 3B). Increasing the dose did not lead to major improvements. Conjugating stP5 to the TAT element led to a slight improvement in activity with 1 μM TAT-stP5 leading to the same 80% reduction visible with 10 μM TAT-P5, likely due to improved cell permeability compared to stP5. Those concentrations are 3 to 10-fold below the K_D_ of stP5 against GEF-H1 measured by SPR (34 μM). The observed bioactivity differences probably reflect the artificial nature of our recombinant DH-PHGEF-H1 protein compared to the full length tightly regulated native version present in cells. While stapled peptides are often characterized by helicity, the helicity degree does not necessarily correlate with improved activity and it is not surprising for such a short sequence like stP5 to not display secondary structure in CD^[47]^.

**Figure 3.**
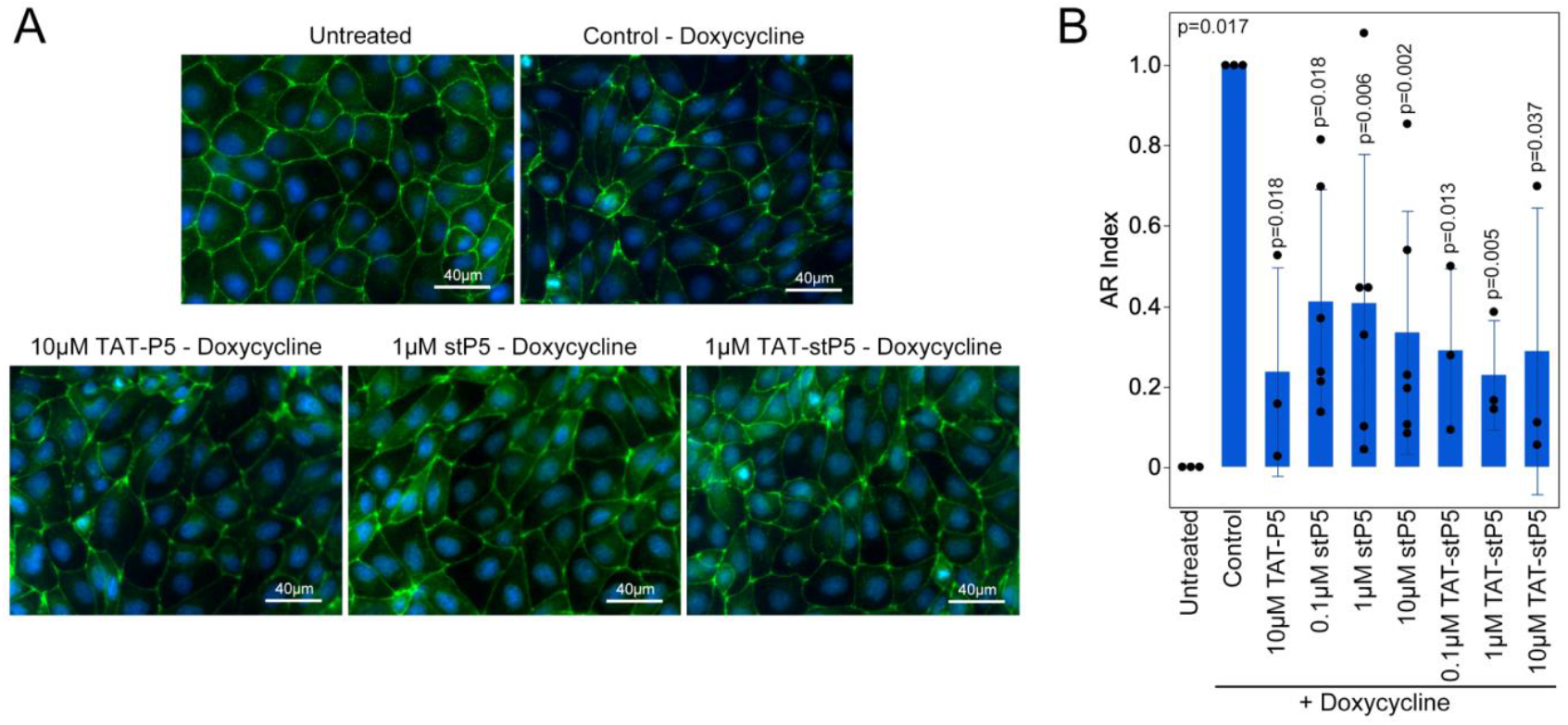
GEF-H1 peptide inhibitors attenuate effects of GEF-H1 overexpression in renal epithelial (MDCK) cells. MDCK/GEF-H1-VSV were left untreated or treated with doxycycline for 24 hours to induce transgene expression in the absence (control) or presence of stP5 or TAT-stP5. The previously characterized TAT-P5 inhibitor was used as a positive control. (**A**) The cells were then fixed and stained for ZO-1 (shown in green, labelling the cell periphery) and DNA (shown in blue). Magnification bar, 40μm. (**B**) To quantify cell shape changes, images were segmented into individual cells and aspect ratios were calculated and used to calculate AR index as a function of the mean AR value and cell-to-cell variability for each analyzed image (AR indices obtained for control samples were used to normalize each experiment). The graph shows quantifications from three independent experiments (shown are means as blue blues with SD and individual data points). Significance was tested with t-tests comparing inhibitor effects to a standard value of 1.

GEF-H1 regulates NF-κB activity in a RhoA-dependent manner and drives expression of pro-inflammatory molecules^[48-49]^. We therefore tested the ability of stP5 to block GEF-H1 induced NF-kB-dependent transcription using a reporter gene assay in cells with GEF-H1-VSV induction by doxycycline and transiently transfected reporter plasmid driving luciferase expression with an NF-κB-dependent promoter (Fig 4A). TAT-P5 led to a dose-dependent reduction of NF-κB promoter activity by 30% at the top concentration tested (20 μM). StP5 was more potent with a dose-dependent reduction of up to 50% at 10 μM. The inhibition of the NF-κB pathway was further improved by improving cell permeability with conjugation of the TAT motif to stP5 (TAT-stP5).

**Figure 4.**
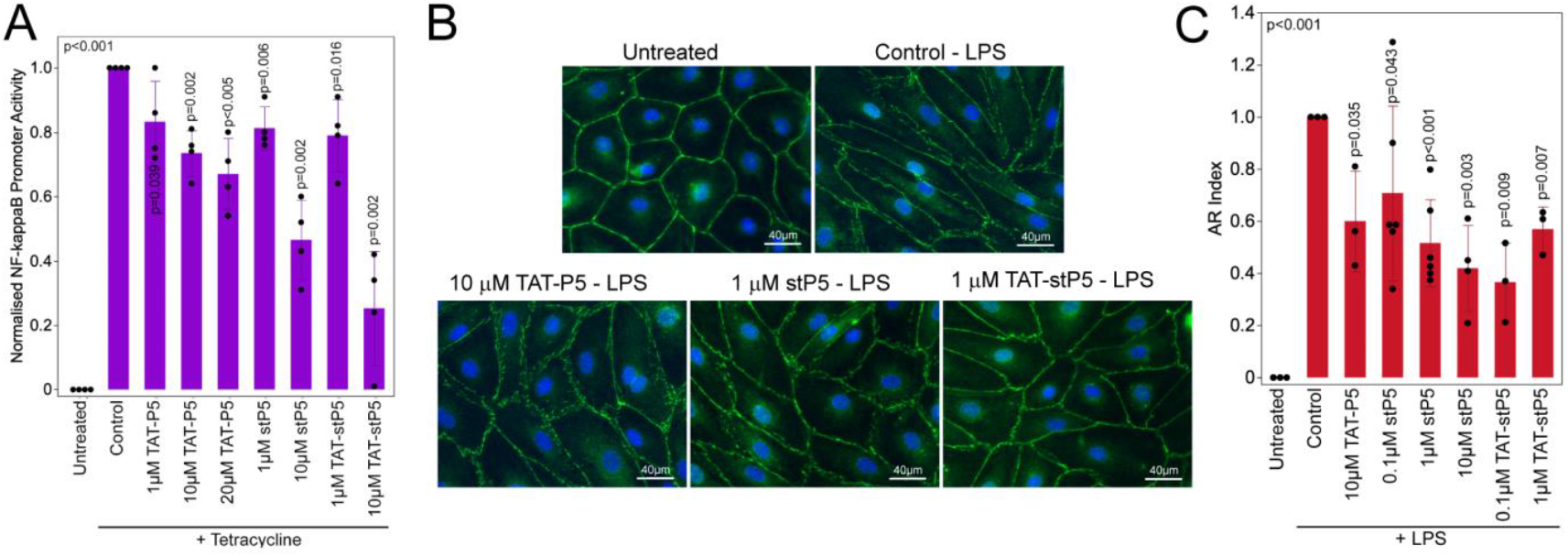
Peptide inhibitors attenuate GEF-H1 induced NF-κB activation and LPS-induced morphological changes in HDMECs. **(A)** MDCK/GEF-H1-VSV cells transiently transfected with reporter plasmids encoding firefly luciferase under the control of an NF-κB-responsive promoter and renilla luciferase under the control of a CMV promoter were incubated for 20 hours with tetracycline prior to cell lysis and measurements of the two luciferase enzyme activities. Firefly to renilla luciferase ratios were calculated and normalized to controls for each experiment. Shown are means of four experiments (bars), standard deviations, and individual data points. Significance was tested with t-tests comparing inhibitor effects to a standard value of HDMECs treated without (control) or with LPS (100ng/ml) in the absence or presence of GEF-H1 peptide inhibitors for 24h. 10 μM TATP5 was used as a positive control. (**B**) Immunofluorescent staining of ZO-1 (Green), which stains the cell periphery, and nuclei (Blue). Magnification bar, 40μm. (**C**) Cell shape changes were quantified by image segmentation and calculation of AR indices as described for MDCK cells in Figure 3. The graph shows quantifications from three independent experiments (shown are means as blue bars with SD and individual data points). Significance was tested with t-tests comparing inhibitor effects to a standard value of 1.

Following the apparent rescue of GEF-H-mediated cellular morphology changes, we wondered if inhibition of GEF-H1/RhoA signaling by stP5 would be useful in pathological contexts. We thus tested stP5 in cellular models of GEF-H1-mediated inflammation. GEF-H1 is activated downstream of numerous cytokines like thrombin and LPS. Cytokine-mediated GEF-H1/RhoA activation in endothelial cells leads to actin remodeling, stress fiber formation and increased cell permeability, ultimately resulting in endothelial barrier dysfunction in lung injury^[31, 50-51]^ and lung infection^[52]^ models. We thus tested the ability of stP5 to prevent LPS-induced cytoskeletal remodeling in human dermal microvascular endothelial cells (HDMECs). Treatments of TAT-P5 (10 μM), stP5 (1 μM) or TAT-stP5 (1 μM) all showed clear partial rescue of the elongated phenotype and junction disruption visible upon treatment of HDMECs with LPS (Fig. 4B). Cells showed overall improved monolayer and intact junction formation. Quantification of cell elongation using the AR index showed that stP5 has dose-dependent and more than 10-fold improved activity compared to TAT-P5, with the 50% rescue observed at 10 μM TAT-P5 recapitulated at 1 μM stP5. Conjugation of the TAT-motif to stP5 led to a further improvement of activity with more than 60% rescue observed at 0.1 μM treatment (Fig. 4C).

StP5 was directly derived from P5, itself designed with a focus on the interaction between GEF-H1 and RhoA specifically^[1]^. These peptides being derived from a nearly native epitope of RhoA, we wondered whether stP5 could inhibit other RhoA-binding GEFs closely related to GEF-H1. This question is further warranted due to the discrepancy between the cell-based activity of around 1 μM and the measured K_D_ against recombinant GEF-H1 (34 μM). p114RhoGEF (or ARHGEF18) is one of the most closely related RhoGEFs to GEF-H1, with 51% identity to the GEF-H1 DHPH domain (SI, Table S1). Spatiotemporal regulation of RhoA by GEF-H1 or p114RhoGEF in epithelial and endothelial cells is a typical example for the specific roles played by individual GEFs. GEF-H1 activity is inhibited by junctional sequestration, in contrast p114RhoGEF is stimulated by cell-cell junction formation and stabilization^[53-55]^. We thus tested TAT-stP5 and stP5 in HDMECs and compared the resulting phenotypes to both GEF-H1 or p114Rho depletion by RNA interference. p114RhoGEF depletion led to marked disruption of junctions. GEF-H1 depletion on the contrary did not affect junctions. Both TAT-stP5 (2 μM) and stP5 (20 μM) did not phenocopy the p114RhoGEF depletion phenotype (Fig. 5A and B). Thus, this sensitive phenotypic readout suggests that TAT-stP5 and stP5 do not inhibit p114RhoGEF; in contrast, they rescue junction disruption induced by activation of GEF-H1 by LPS (Fig 5A and B).

**Figure 5.**
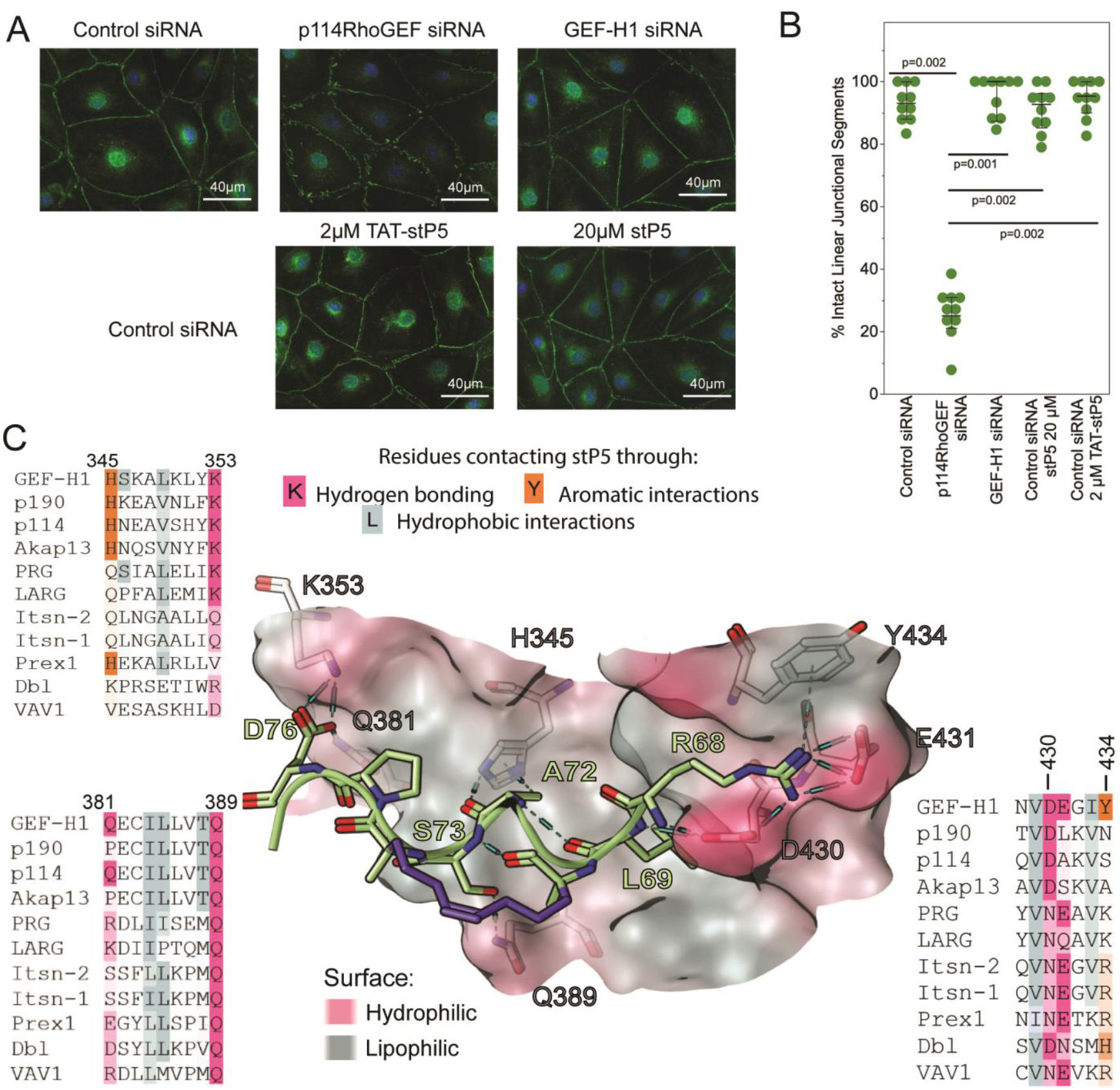
GEF-H1 inhibitors do not phenocopy p114RhoGEF/ARHGEF18 depletion. HDMEC cells were transfected with nontargeting control siRNAs or siRNAs specific for either p114RhoGEF or GEF-H1. Control siRNA transfected cells were then incubated without or with GEF-H1 inhibitors for 24 hours. (**A**) Immunofluorescent staining of the junctional protein ZO-1 (Green) and nuclei (Blue). Note, p114RhoGEF-depletion induces discontinuous ZO-1-positive disrupted tight junctions^[53-54, 58]^ but not depletion of GEF-H1 or either of the GEF-H1 inhibitors (**B**) Images derived from two separate experiments were then quantified by counting intact linear junctional segments between two cells and expressing the values as a percentage of the total junctional segments visible in an image (shown are means as green bars with SD and individual data points). A Steel-Dwaas test was used to calculate p-values. Note, higher concentrations of GEF-H1 inhibitors were tested than in the experiments testing inhibition of GEF-H1. (**C**) Analysis of the binding site of stP5 in a GEF-H1 model in a bioinformatic context. The peptide is in light green, with its staple in purple. The 10 most related RhoGEFs by DHPH sequence are aligned in the regions interacting with the peptide. Residues of GEF-H1 undergoing hydrogen bonds (via side-chains) are highlighted in pink, pi-stacking interactions in orange, and primarily hydrophobic interactions in blue. Residue numbering corresponds to GEF-H1. The corresponding residues in other GEFs are highlighted in the same colour, fading with lowering similarity to GEF-H1. A full alignment with all binding site residues can be found in the SI.

We then investigated whether stP5 selectivity for GEF-H1 could result from structural changes in GEF bindings sites, and if these structural changes could be harnessed for future improved designs. The binding mode of stP5 in GEF-H1 was modeled by energy minimization in a rigid GEF-H1 homology model, based on where the natural RhoA-GEF interaction takes place. The theoretical binding sites of related RhoGEFs were compared by sequence (Fig. 5C). A full sequence alignment of the DHPH domains highlighting binding site residues can be found in SI Fig. S3. Key interactions occur via stP5 residues R68, L69, A72, D76 on one helix face and via S73 on the other face. Residue L69 buries in the deepest part of the binding site, surrounded by a conserved hydrophobic patch across RhoGEFs. L69 has been previously identified as a recurrent hot-spot residue at the interface between RhoGTPases and GEFs^[56]^, as per alanine scanning mutagenesis showing a ΔΔG of 2.4-2.9 kcal/mol for all related RhoGEF-GTPase complexes investigated (SI, Table S2) indicating conserved importance of L69. Residue A72, originally L72 in Rho, is mutated in stP5 to avoid clashes with GEF-H1 residue H345^[1]^. A72 is predicted to undergo a CH_3_-π stacking interaction with the imidazole ring of H345, and the backbone of stP5 contacts the imidazole N-H through H-bonding. H345 is conserved in the closest GEFs to GEF-H1, but more distant GEFs like Dbl do not bear aromatic residues there. S73 undertakes an H-bond with highly conserved residue Q389 and is thus unlikely to discriminate between RhoGEFs. D76 is an interesting residue as it was previously shown to govern the selectivity of RhoGEFs for certain RhoGTPases^[39, 57]^. D76 is, in fact, the only amino acid in stP5 differing within the sequence of Rho family members, mutated to Q in Cdc42 and Rac1 (SI. Fig. S4).

D76 contacts GEF-H1 through a salt bridge with K353 and Q381. Remarkably, K353 is conserved across GEFs known to exchange RhoA (top 5 GEFs), whereas GEFs exchanging Cdc42 or Rac1 like Intersectins (Itsns) and Prex1 do not bear residues capable of salt bridges at this position, providing some basis for selectivity determination by D76. On the other hand, Q381 is considerably less conserved within RhoA-exchanging GEFs. The alanine scanning mutagenesis energy of D76 in our GEF-H1/RhoA model was compared to available related DHPH-Rho crystal structures in the PDB (Table 1), revealing that the dual presence of Q381 and K353 in GEF-H1 (and p114RhoGEF) considerably increases the ΔΔG of D76 compared to all GEFs investigated. Taken together, A72 and D76 can plausibly account for some of the observed stP5’s affinity for GEF-H1 over non-RhoA exchanging GEFs.

**Table 1.**
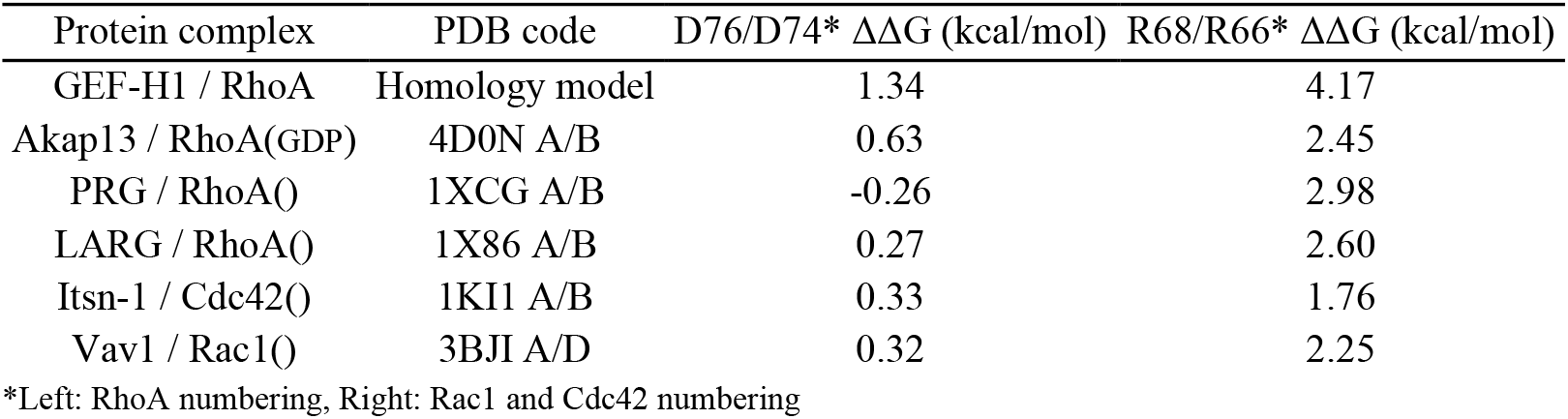
Alanine Scanning mutagenesis results (Robetta software) for Rho residues D76 and R68 in complex with GEFs.

The last residue in stP5, R68, could, in turn, be the main reason for stP5’s preferential inhibition of GEF-H1 over p114RhoGEF. R68 is involved in numerous polar interactions with a non-conserved polar interface in GEF-H1 in our model, consisting of multiple H-bonds taking place between the guanidine moiety and E431 and D430, as well as a potential π-stacking interaction with Y434. Firstly, amino acid D430 is mutated to N in more distant GEFs, likely affecting the strength of the H-bond occurring with the R68 backbone. For more closely related GEFs like p114RhoGEF, amino acids E431 and Y343 may be important as p114RhoGEF bears a lipophilic leucine at position 431 and non-aromatic residue asparagine at position 434, both lacking the necessary properties to reproduce the same R68 binding mode as GEF-H1. To further support the importance of R68 as a GEF-H1-specific hot-spot residue, alanine scanning mutagenesis energy of R68 in the GEF-H1-RhoA model shows a dramatic increase to 4.2 kcal/mol compared to related DHPH-Rho crystal structures (Table 1). In conclusion, molecular modelling and alanine scanning mutagenesis suggest that residues D76 and R68 in stP5 may govern preferential inhibition for GEF-H1 over p114RhoGEF.

In summary, we report that short and stapled peptide stP5 (10 residues) as a GEF-H1 inhibitor with improved properties compared to our previously reported peptide TAT-P5 (31 aa)^[1]^. The molecule was derived using a ligand-based drug design approach by truncating and stapling the helical core of the RhoA epitope P5. StP5 shows equilibrium binding to GEF-H1 by SPR and rescues morphological changes induced by GEF-H1 overexpression in MDCK cells. Critically the stapling modification resulted in around a 100 fold improvement in activity in the cell-based assays. We then show that GEF-H1 inhibition by stP5 is a viable approach to rescue transcriptional activation of NF-κB reporter and cytoskeletal rearrangements induced by LPS in endothelial cells. Thus, these studies show that GEF-H1 inhibition could be a viable approach to tackle inflammatory disorders affecting the endothelium.

StP5 further showed selectivity for GEF-H1 compared to a closely related p114RhoGEF, as measured by a robust phenotypic assay. A bioinformatic analysis of the stP5 binding mode in GEF-H1 suggests that stP5 may have differential binding affinities to different GEFs mediated by specific hot-spots identified in the stP5 sequence by *in silico* alanine scanning mutagenesis experiments. While this is only preliminary evidence for the selectivity of stP5 for GEF-H1, and binding affinity may only be one of the facets affecting stP5s activity in cells, this initial result is encouraging for further design of GEF-specific inhibitors.

Targeting GTPase signaling through GEFs is an emerging strategy which requires novel drug discovery approaches to address the challenges associated with the target features involved. GEF-H1 has been identified as overexpressed in patient tissues suffering from different carcinoma^[34-35, 59]^, melanoma^[32]^ and fibrotic diseases^[29]^ by exacerbating Rho signaling. Our developed peptide stP5 will thus contribute to the validation of GEF-H1 as a druggable target in such disorders. We also anticipate that the bioinformatic study described herein, pinning down key hot spots from the GEF-H1/RhoA interaction, will demonstrate the feasibility of the GEF-targeting approach and pave the way for improved peptidic or small molecule designs.

Limitations of our study. Our peptide stP5 demonstrates higher cellular potency and greater selectivity than might be expected from the in vitro binding data. Readily observable helicity was not detected by CD though could be induced by binding to GEF-H1. Further investigations into stabilized peptides targeting GEF-H1 will be required to understand fully these aspects.

## AUTHOR INFORMATION

### Corresponding Authors

D. Selwood and M. S. Balda

### Author Contributions

EC, DLS, MSB and KM conceived the study. EC designed stP5 and supervised molecular modelling experiments performed by CG. MSB and KM supervised cell culture experiments performed by TL. DLS designed and supervised synthesis, protein purification and SPR experiments performed by CG. WCW and SY performed the CD. CG, EC, DLS, MSB and KM wrote the manuscript.

## ABBREVIATIONS

PPI: Protein-protein interaction
GEF: Guanine exchange factor
GDP: guanine diphosphate
GTP: guanine triphosphate
RhoA: Ras homolog A
, DH: Dbl homology
PH: pleckstrin homology
LPS: lipopolysaccharides, twin-arginine translocation - TAT
PAIN: Pan-assay interference
SPR: surface plasmon resonance
MDCK: Madin-Darby Canine Kidney cells
αSMA: α-smooth muscle actin
SRE: serum response elements
LDH: lactate dehydrogenase
TGF-β: Transforming Growth Factor β
HDMECs: Human Dermal Micorvascular Endothelial Cells
H-bond: Hydrogen bond
Itsn: Interesectin
Cdc42: Cell division control protein 42 homolog
AKAP13: A-kinase anchoring protein 13 – Akap13
PRG: PDZ-RhoGEF
LARG: Leukemia-associated RhoGEF
Prex1: phosphatidylinositol-3,4,5-trisphosphate dependent Rac exchange factor 1
vav1: vav guanine nucleotide exchange factor 1

