## Supplementary information for "Inhibition of GEF-H1-RhoA signaling in inflammation with a stapled peptide mimicry of the RhoA^67-78^ helix"

### Supporting information

#### Supplementary figures and tables

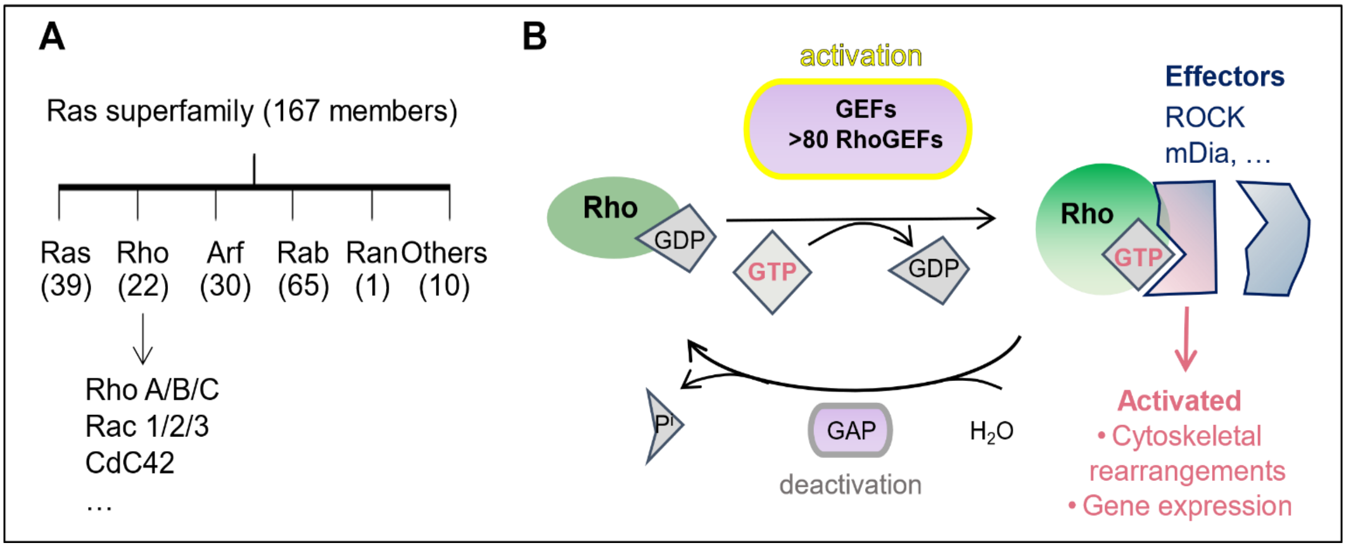

Figure S1. The Ras GTPases superfamily structure and principle of the GTPase switch – **A**. The Rho subfamily comprises about 22 members, the most studied being RhoA, Cdc42 and Rac1. **B**. RhoA is inactive when bound to GDP. Upon exchange for GTP, RhoA undergoes conformational changes increasing its affinity for effectors and triggers cellular processes upon binding them. Nucleotide exchange is catalyzed by GEFs, whereas GAPs catalyze GTP hydrolysis to inactivate GTPases.

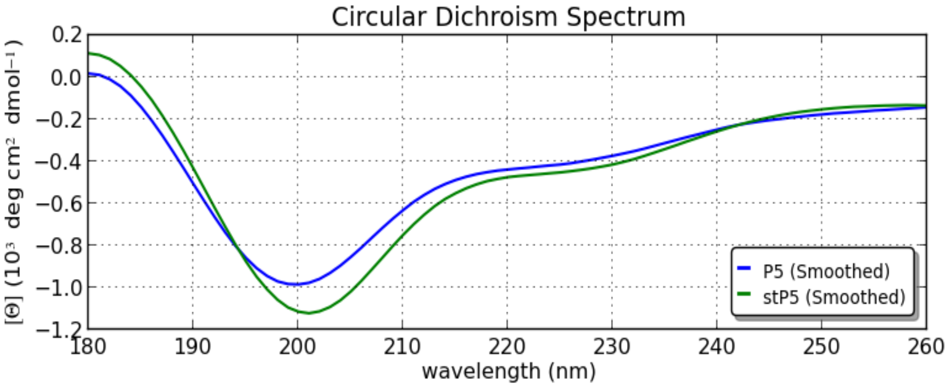

Figure S2. Circular Dichroism spectrum to compare the helicity of stP5 and P5.

The initial concentration of both samples was 125 µM and was then diluted to 62.5 µM with 50% of trifluoroethanol. Each line represents the mean of 10 repeated scans with a 1 nm bandwidth and a time-per-point of 1 s for a nominal step size of 1.0 nm. Data analysis was performed using the BeStSel website[1].

#### Table S1. Similarity matrix for related GEFs to the DHPH domain of GEF-H1 (Clustalω)

| | | | | | | | | | | | |
|---|---|---|---|---|---|---|---|---|---|---|---|
| 1: sp\|Q92974\|GEF-H1 | 100.00 | 53.13 | 50.74 | 50.15 | 31.00 | 28.48 | 25.79 | 24.92 | 21.43 | 21.38 | 21.00 |
| 2: sp\|Q8N1W1\|p190 | 53.13 | 100.00 | 39.11 | 31.31 | 20.05 | 18.11 | 17.60 | 17.53 | 16.34 | 18.69 | 18.35 |
| 3: sp\|Q6ZSZ5\|p114 | 50.74 | 39.11 | 100.00 | 36.95 | 22.77 | 20.66 | 18.29 | 16.50 | 18.45 | 16.12 | 17.79 |
| 4: sp\|Q12802\|Akap13 | 50.15 | 31.31 | 36.95 | 100.00 | 21.10 | 18.03 | 17.00 | 16.64 | 14.83 | 16.95 | 15.84 |
| 5: sp\|O15085\|PRG | 31.00 | 20.05 | 22.77 | 21.10 | 100.00 | 37.52 | 17.85 | 17.53 | 19.22 | 18.57 | 17.93 |
| 6: sp\|Q9NZN5\|LARG | 28.48 | 18.11 | 20.66 | 18.03 | 37.52 | 100.00 | 17.41 | 17.44 | 18.76 | 17.12 | 19.05 |
| 7: sp\|Q9NZM3\|Itsn-2 | 25.79 | 17.60 | 18.29 | 17.00 | 17.85 | 17.41 | 100.00 | 62.49 | 19.21 | 15.04 | 15.90 |
| 8: sp\|Q15811\|Itsn-1 | 24.92 | 17.53 | 16.50 | 16.64 | 17.53 | 17.44 | 62.49 | 100.00 | 19.58 | 15.36 | 16.54 |
| 9: sp\|Q8TCU6\|Prex1 | 21.43 | 16.34 | 18.45 | 14.83 | 19.22 | 18.76 | 19.21 | 19.58 | 100.00 | 21.88 | 22.04 |
| 10: sp\|P10911\|Dbl | 21.38 | 18.69 | 16.12 | 16.95 | 18.57 | 17.12 | 15.04 | 15.36 | 21.88 | 100.00 | 19.08 |
| 11: sp\|P15498\|Vav1 | 21.00 | 18.35 | 17.79 | 15.84 | 17.93 | 19.05 | 15.90 | 16.54 | 22.04 | 19.08 | 100.00 |

```
GEF-H1  ---------------------------------KQQDVIYELIQTELHHVRTLKIMTRLF 261
p190    EAES---------WSLVVDPS-F-CNRQEKDVIKRQDVIFELMQTEMHHIQTLFIMSEIF 875
p114    EAES---------WSLAVDAA-Y-AKKQKREVVKRQDVLYELMQTEVHHVRTLKIMLKVY 473
Akap13  EAES---------WSRIIDSK-F-LKQQKKDVVKRQEVIYELMQTEFHHVRTLKIMSGVY 2020
PRG     DAQN---------WQHTVGKD-V-VAGLTQREIDRQEVINELFVTEASHLRTLRVLDLIF 760
LARG    DPPN---------WQQLVSRE-V-LLGLKPCEIKRQEVINELFYTERAHVRTLKVLDQVF 813
Itsn-2  -------------WCADL----QTLDTMQPIERKRQGYIHELIQTEERYMADLQLVVEVF 1235
Itsn-1  -------------WCSDL----HLLDMLTPTERKRQGYIHELIVTEENYVNDLQLVTEIF 1263
Prex1   CAHPDPRAPGAAAPSSGPGPCAA-ARESERQLRLRLCVLNEILGTERDYVGTLRFLQSAF 75
Dbl     CQEK---------RSSGPSSS-L-DNG-NSLDVLKNHVLNELIQTERVYVRELYTVLLGY 521
VAV1    CVEN-EEAEGDEIYEDLMRSEPVSMPPKMTEYDKRCCCLREIQQTEEKYTDTLGSIQQHF 2020
                                          :   : *:  **  :   *  :   :
                                                     DH DOMAIN

GEF-H1  RTGMLEELHLE------PGVVQGLFPCVDELSDIHTRFLSQLLERRRQALCPGSTRNFVI 315
p190    RKGMKEELQLD------HSTVDKIFPCLDELLEIHRHFFYSMKERRQE-SCAGSDRNFVI 928
p114    SRALQEELQFS------SKAIGRLFPCADDLLETHSHFLARLKERRQESLEEGSDRNYVI 527
Akap13  SQGMMADLLFE------QQMVEKLFPCLDELISIHSQFFQRILERKKESLVDKSEKNFLI 2074
PRG     YQRMKKENLMP------REELARLFPNLPELIEIHNSWCEAMKKLREE--------GPII 806
LARG    YQRVSREGILS------PSELRKIFSNLEDILQLHIGLNEQMKAVRKR------NETSVI 861
Itsn-2  QKRMAESGFLT------EGEMALIFVNWKELIMSNTKLLKALRVRKKTG-----GEKMPV 1284
Itsn-1  QKPLMESELLT------EKEVAMIFVNWKELIMCNIKLLKALRVRKKMS-----GEKMPV 1312
Prex1   LHRIRQNVADSVEKGLTEENVKVLFSNIEDILEVHKDFLAALEY-CLHP------EPQSQ 128
Dbl     RAEMDNPEMFDLMPPLLRNKKDILFGNMAEIYEFHNDIFLSSLENCAH----------AP 571
VAV1    LKPLQR--------FLKPQDIEIIFINIEDLLRVHTHFLKEMKEALGTP---G------A 263
           :                   :*    ::   :
                                DH DOMAIN

GEF-H1  HRLGDLLISQFSGPSAEQMCKTYSEFCSRHSKALKLYKELYARDKRFQQFIRKVTR--PA 373
p190    DRIGDILVQQFSEENASKMKKIYGEFCCHHKEAVNLFKEL-QQNKKFQNFIKLRNS--NL 985
p114    QKIGDLLVQQFSGENGERMKEKYGVFCSGHNEAVSHYKLLLQQNKKFQNLIKKIGN--FS 585
Akap13  KRIGDVLVNQFSGENAERLKKTYGKFCGQHNQSVNYFKDLYAKDKRFQAFVKKKMS--SS 2132
PRG     KEISDLMLARFDGPAREELQQVAAQFCSYQSIALELIKTKQRKESRFQLFMQEAES--HP 864
LARG    DQIGEDLLTWFSGPGEEKLKHAAATFCSNQPFALEMIKSRQKKDSRFQTFVQDAES--NP 919
Itsn-2  QMIGDILAAE------LSHMQAYIRFCSCQLNGAALLQQKTDEDTDFKEFLKKLAS--DP 1336
Itsn-1  KMIGDILSAQ------LPHMQPYIRFCSRQLNGAALIQQKTDEAPDFKEFVKRLAM--DP 1364
Prex1   HELGNVFLKF------KDKFCVYEEYCSNHEKALRLLVELNK-IPTVRAFLLSCMLLGGR 181
Dbl     ERVGPCFLER------KDDFQMYAKYCQNKPRSETIWRK-YSECAFFQECQ-------RK 617
VAV1    ANLYQVFIKY------KERFLVYGRYCSQVESASKHLDRVAAAREDVQMKLEECSQ--RA 315
          :   :                  :*     .             .:
                                DH DOMAIN

GEF-H1  VLKRHGVQECILLVTQRITKYPLLISRILQHSHGIEEERQDLTTALGLVKELLSNVDEGI 433
p190    LARRRGIPECILLVTQRITKYPVLVERILQYTKERTEEHKDLRKALCLIKDMIATVDLKV 1045
p114    IVRRLGVQECILLVTQRITKYPVLVERIIQNTEAGTEDYEDLTQALNLIKDIISQVDAKV 645
Akap13  VVRRLGIPECILLVTQRITKYPVLFQRILQCTKDNEVEQEDLAQSLSLVKDVIGAVDSKV 2192
PRG     QCRRLQLRDLIISEMQRLTKYPLLLESIIKHTEGGTSEHEKLCRARDQCREILKYVNEAV 924
LARG    LCRRLQLKDIIPTQMQRLTKYPLLLDNIAKYTEWP-TEREKVKKAADHCRQILNYVNQAV 978
Itsn-2  RCKGMPLSSFLLKPMQRITRYPLLIRSILENTPESHADHSSLKLALERAEELCSQVNEGV 1396
Itsn-1  RCKGMPLSSFILKPMQRVTRYPLIIKNILENTPENHPDHSHLKHALEKAEELCSQVNEGV 1424
Prex1   KTTDIPLEGYLLSPIQRICKYPLLLKELAKRTPGKHPDHPAVQSALQAMKTVCSNINETK 241
Dbl     LKHRLRLDSYLLKPVQRITKYQLLLKELLKYSKDC-EGSALLKKALDAMLDLLKSVNDSM 676
VAV1    NNGRFTLRDLLMVPMQRVLKYHLLLQELVKHTQEA-MEKENLRLALDAMRDLAQCVNEVK 374
              :   :    **: :* ::.  : : :         :  :      :   ::
                                DH DOMAIN
```

Figure S3 – Sequence alignment for the 10 closest related GEFs to the GEF-H1 DHPH domain by sequence (BLAST). Consensus is shown at the bottom. Coloring corresponds to the stP5 binding site, coded according to the stP5 residue near the GEF-H1 residue.

```
GEF-H1   YQLEKGARLQEIYNRMDPRAQTPVPGKGPFGREELLR--RKLIHDGCLLWKTA---TGRF 502
p190     NEYEKNQKWLEILNKIENKTYTKLKNGHVFRKQALMSEERTLLYDGLVYWKTA---TGRF 1116
p114     SECEKGQRLREIAGKMDLKSSSKLKNGLTFRKEDMLQ--RQLHLEGMLCWKTT---SGRL 714
Akap13   ASYEKKVRLNEIYTKTDSKSIMRMKSGQMFAKEDLKR--KKLVRDGSVFLKNA---AGRL 2261
PRG      KQTENRHRLEGYQKRLDATALERASNPLAAEFKSLDLTTRKMIHEGPLTWRIS---KDKT 995
LARG     KEAENKQRLEDYQRRLDTSSLKLSEYPNVEELRNLDLTKRKMIHEGPLVWKVN---RDKT 1049
Itsn-2   REKENSDRLEWIQAHVQCEGLAE---QLIFNSLTNCLGPRKLLHSGKLYK------TKSN 1461
Itsn-1   REKENSDRLEWIQAHVQCEGLSE---QLVFNSVTNCLGPRKFLHSGKLYK------AKSN 1489
Prex1    RQMEKLEALEQLQSHIEGWEGSN-----------LTDICTQLLLQGTLL-KIS---AGNI 307
Dbl      HQIA----INGYIGNLNELG-------------------KMIMQGGFSVWIGHKKGATKM 735
VAV1     RDNETLRQITNFQLSIENLDQSLAHYGRPKIDGELKI------------TSVE---RRSK 433
          .               :
```

```
                                                          PH DOMAIN
GEF-H1   KDVLVLLMTDVLVFLQEKDQKYIFPTLD--------------KPSVVSLQNLIVRDIAN 543
p190     KDILALLLTDVLLFLQEKDQKYIFAAVD--------------QKPSVISLQKLIAREVAN 1148
p114     KDILAILLTDVLLLLQEKDQKYVFASVD--------------SKPPVISLQKLIVREVAN 746
Akap13   KEVQAVLLTDILVFLQEKDQKYIFASLD--------------QKSTVISLKKLIVREVAH 2293
PRG      LDLHVLLLEDLLVLLQKQDEKLLLKCHSKTAVGSS---DSKQTFSPVLKLNAVLIRSVAT 1038
LARG     IDLYTLLLEDILVLLQKQDDRLVLRCHSKILASTA---DSKHTFSPVIKLSTVLVRQVAT 1092
Itsn-2   KELHGFLFNDFLLLTYMVKQFAVSSGSEKLFSSKSNAQFKMYKTPIFLNEVLVKLPTDPS 1505
Itsn-1   KELYGFLFNDFLLLTQITKPLG-SSGTDKVFSPKSNLQYKMYKTPIFLNEVLVKLPTDPS 1532
Prex1    QERAFFLFDNLLVYCKRKSRVTGS---KKSTKRTKSINGSLYIFRGRINTEVMEVENVED 351
Dbl      KDLARFKPMQRHLFLYEKAI---VFCKRRVESGEGSDRYPSYSFKHCWKMDEVGITEYVK 770
VAV1     MDRYAFLLDKALLICKRRGDSY-------------------DLKDFVNLHSFQVRDDSS 477
          :   .   .  :                                .   .
```

```
                                PH DOMAIN
GEF-H1   QEK----------GMFLISA---APPEMYEVHTASRDDRSTWIRVIQQSVR--------- 571
p190     EER----------GMFLISAS-SAGPEMYEIHTNSKEERNNWMRRIQQAVESCP-EEKGG 1196
p114     EEK----------AMFLISAS-LQGPEMYEIYTSSKEDRNAWMAHIQRAVESCP-DEEEG 794
Akap13   EEK----------GLFLISMG-MTDPEMVEVHASSKEERNSWIQIIQDTINTLNRDEDEG 2342
PRG      DKR----------AFFIICTSKLGPPQIYELVALTSSDKNTWMELLEEAVRNATRHPGAA 1088
LARG     DNK----------ALFVISMS-DNGAQIYELVAQTVSEKTVWQDLICRMAASVK-EQSTK 1140
Itsn-2   SDE----------PVFHISHI----DRVYTLRTDNINERTAWVQKIKAASEQYIDTEKKK 1552
Itsn-1   GDE----------PIFHISHI----DRVYTLRAESINERTAWVQKIKAASELYIETEKKK 1580
Prex1    GTADYHSNGYTVTNGWKIHNT--AKNKWFVCMAKTAEEKQKWLDAIIREREQRESLKLGM 401
Dbl      GDN----------RKFEIWYG--EKEEVYIVQASNVDVKMTWLKEIRNILLKQQEL---- 814
VAV1     GDRDNKK----WSHMFLLIED--QGAQGYELFFKTRELKKKWMEQFEMAISNIYPENATA 513
                                PH DOMAIN
```

Region surrounding D76 | Region surrounding A72 and S73
Region surrounding L69 | Region surrounding R68

Figure S3 (cont.) – Sequence alignment for the 10 closest related GEFs to the GEF-H1 DHPH domain by sequence (BLAST). Consensus is shown at the bottom. Coloring corresponds to the stP5 binding site, coded according to the stP5 residue near the GEF-H1 residue.

```
                              Switch I                Switch II     P5
                                                                    stP5
RhoA    CGKTCLLIVFSKDQFPEVYVPTVFENYVADIEVDGKQVELALWDTA GQEDYDRLRPLSYPDTDVILMCFSIDSPDSLENIPEKWT  100
Rac1    VGKTCLLISYTTNAFPGEYIPTVFDNYSANVMVDGKPVNLGLWDTA GQEDYDRLRPLSYPQTDVFLICFSLVSPASFENVRAKWY  98
Cdc42   VGKTCLLISYTTNKFPSEYVPTVFDNYAVTVMIGGEPYTLGLFDTA GQEDYDRLRPLSYPQTDVFLVCFSVVSPSSFENVKEKWV  98
        ******* :::.: **  *:****:** : : :.*:   *:*:*****************:***:*:***: ** *:**:  **
```

Figure S4 – Sequence alignment of RhoA, Rac1 and Cdc42, the three main members of the Rho GTPase family. Consensus is shown at the bottom. Coloring corresponds to the stP5 and P5 regions.

Table S2. Extended data: Alanine scanning mutagenesis generated with Robetta software raw data

Results were generated with the Robetta software with PDB files as the input. Rho residues at the interface with GEFs investigated in this study are highlighted.

**GEF-H1**

| pdb | #chain | int_id | res# | aa | DDG (complex) | DDG (complex,obs) | DG (partner) | |
|---|---|---|---|---|---|---|---|---|
| 63 | B | 1 | 429 | 14 | 0.50 | 0.00 | 0.90 | Q |
| 64 | B | 0 | 430 | 4 | -0.09 | 0.00 | 2.75 | E |
| 65 | B | 1 | 431 | 3 | 0.68 | 0.00 | -0.94 | D |
| 66 | B | 1 | 432 | 20 | 2.60 | 0.00 | 0.40 | Y |
| 67 | B | 0 | 433 | 3 | -0.02 | 0.00 | 0.45 | D |
| 68 | B | 1 | 434 | 15 | 4.17 | 0.00 | -0.03 | R |
| 69 | B | 1 | 435 | 10 | 2.64 | 0.00 | 0.64 | L |
| 70 | B | 0 | 436 | 15 | 0.00 | 0.00 | 4.54 | R |
| 72 | B | 1 | 438 | 10 | 2.02 | 0.00 | 0.84 | L |
| 73 | B | 1 | 439 | 16 | 1.08 | 0.00 | 0.07 | S |
| 74 | B | 0 | 440 | 20 | 0.00 | 0.00 | 3.09 | Y |
| 76 | B | 1 | 442 | 3 | 1.34 | 0.00 | -0.38 | D |
| 77 | B | 0 | 443 | 17 | 0.00 | 0.00 | 2.71 | T |
| 78 | B | 0 | 444 | 3 | 0.00 | 0.00 | 0.15 | D |
| 79 | B | 0 | 445 | 18 | 0.00 | 0.00 | 2.43 | V |
| 80 | B | 0 | 446 | 8 | 0.00 | 0.00 | 3.48 | I |

**Akap13 (4D0N)**

| pdb | #chain | int_id | res# | aa | DDG (complex) | DDG (complex,obs) | DG (partner) | |
|---|---|---|---|---|---|---|---|---|
| 63 | A | 1 | 61 | 14 | 1.90 | 0.00 | 1.52 | Q |
| 64 | A | 0 | 62 | 4 | -0.05 | 0.00 | 4.16 | E |
| 65 | A | 0 | 63 | 3 | -0.04 | 0.00 | -0.74 | D |
| 66 | A | 1 | 64 | 20 | 2.09 | 0.00 | 0.30 | Y |
| 67 | A | 0 | 65 | 3 | -0.03 | 0.00 | 0.21 | D |
| 68 | A | 1 | 66 | 15 | 2.45 | 0.00 | -0.26 | R |
| 69 | A | 1 | 67 | 10 | 2.45 | 0.00 | 0.73 | L |
| 70 | A | 0 | 68 | 15 | -0.01 | 0.00 | 4.48 | R |
| 72 | A | 1 | 70 | 10 | 1.93 | 0.00 | 0.81 | L |
| 73 | A | 1 | 71 | 16 | 3.02 | 0.00 | 0.46 | S |
| 74 | A | 0 | 72 | 20 | 0.00 | 0.00 | 3.14 | Y |
| 76 | A | 1 | 74 | 3 | 0.63 | 0.00 | -0.37 | D |
| 77 | A | 0 | 75 | 17 | 0.00 | 0.00 | 1.13 | T |
| 78 | A | 0 | 76 | 3 | 0.00 | 0.00 | 0.01 | D |
| 79 | A | 0 | 77 | 18 | 0.00 | 0.00 | 2.49 | V |
| 80 | A | 0 | 78 | 8 | 0.00 | 0.00 | 3.67 | I |

**PRG (1XCG)**

| pdb | #chain | int_id | res# | aa | DDG (complex) | DDG (complex,obs) | DG (partner) | |
|---|---|---|---|---|---|---|---|---|
| 63 | B | 1 | 416 | 14 | 0.37 | 0.00 | 1.13 | Q |
| 64 | B | 0 | 417 | 4 | -0.10 | 0.00 | 2.95 | E |
| 65 | B | 0 | 418 | 3 | -0.06 | 0.00 | -0.77 | D |
| 66 | B | 1 | 419 | 20 | 2.81 | 0.00 | 0.21 | Y |
| 67 | B | 0 | 420 | 3 | -0.04 | 0.00 | 0.06 | D |
| 68 | B | 1 | 421 | 15 | 2.98 | 0.00 | 0.01 | R |
| 69 | B | 1 | 422 | 10 | 2.85 | 0.00 | 0.60 | L |
| 70 | B | 0 | 423 | 15 | 0.01 | 0.00 | 3.89 | R |
| 72 | B | 1 | 425 | 10 | 1.42 | 0.00 | 0.62 | L |
| 73 | B | 0 | 426 | 16 | -0.05 | 0.00 | 0.28 | S |
| 74 | B | 0 | 427 | 20 | 0.00 | 0.00 | 3.12 | Y |
| 76 | B | 1 | 429 | 3 | -0.26 | 0.00 | -0.44 | D |
| 77 | B | 0 | 430 | 17 | 0.00 | 0.00 | 1.31 | T |
| 78 | B | 0 | 431 | 3 | 0.00 | 0.00 | -0.46 | D |
| 79 | B | 0 | 432 | 18 | 0.00 | 0.00 | 2.37 | V |
| 80 | B | 0 | 433 | 8 | 0.00 | 0.00 | 3.82 | I |

**LARG (1X86)**

| pdb | #chain | int_id | res# | aa | DDG (complex) | DDG (complex,obs) | DG (partner) | |
|---|---|---|---|---|---|---|---|---|
| 63 | B | 1 | 424 | 14 | 0.33 | 0.00 | 1.03 | Q |
| 64 | B | 0 | 425 | 4 | -0.04 | 0.00 | 3.08 | E |
| 65 | B | 0 | 426 | 3 | -0.04 | 0.00 | -0.56 | D |
| 66 | B | 1 | 427 | 20 | 2.58 | 0.00 | 0.40 | Y |
| 67 | B | 0 | 428 | 3 | -0.02 | 0.00 | -0.49 | D |
| 68 | B | 1 | 429 | 15 | 2.60 | 0.00 | -0.24 | R |
| 69 | B | 1 | 430 | 10 | 2.39 | 0.00 | 0.59 | L |
| 70 | B | 0 | 431 | 15 | 0.00 | 0.00 | 3.48 | R |
| 72 | B | 1 | 433 | 10 | 1.80 | 0.00 | 0.50 | L |
| 73 | B | 0 | 434 | 16 | -0.02 | 0.00 | -0.22 | S |
| 74 | B | 0 | 435 | 20 | 0.00 | 0.00 | 3.25 | Y |
| 76 | B | 1 | 437 | 3 | 0.27 | 0.00 | -0.44 | D |
| 77 | B | 0 | 438 | 17 | -0.01 | 0.00 | 0.75 | T |
| 78 | B | 0 | 439 | 3 | -0.04 | 0.00 | -0.08 | D |
| 79 | B | 0 | 440 | 18 | 0.00 | 0.00 | 1.91 | V |
| 80 | B | 0 | 441 | 8 | 0.00 | 0.00 | 3.48 | I |

**Itsn-1 (1KI1)**

| pdb | #chain | int_id | res# | aa | DDG (complex) | DDG (complex,obs) | DG (partner) | |
|---|---|---|---|---|---|---|---|---|
| 61 | A | 1 | 61 | 14 | 1.13 | 0.00 | 0.83 | Q |
| 62 | A | 0 | 62 | 4 | -0.08 | 0.00 | 4.04 | E |
| 63 | A | 0 | 63 | 3 | -0.01 | 0.00 | -0.64 | D |
| 64 | A | 1 | 64 | 20 | 1.87 | 0.00 | 0.64 | Y |
| 65 | A | 1 | 65 | 3 | 1.42 | 0.00 | -1.14 | D |
| 66 | A | 1 | 66 | 15 | 1.76 | 0.00 | -0.45 | R |
| 67 | A | 1 | 67 | 10 | 2.54 | 0.00 | 0.85 | L |
| 68 | A | 0 | 68 | 15 | 0.00 | 0.00 | 2.24 | R |
| 70 | A | 1 | 70 | 10 | 1.82 | 0.00 | 0.85 | L |
| 71 | A | 1 | 71 | 16 | 0.65 | 0.00 | 0.11 | S |
| 72 | A | 0 | 72 | 20 | 0.00 | 0.00 | 4.26 | Y |
| 74 | A | 1 | 74 | 14 | 0.33 | 0.00 | -0.55 | D |
| 75 | A | 0 | 75 | 17 | 0.00 | 0.00 | 1.16 | T |
| 76 | A | 0 | 76 | 3 | 0.00 | 0.00 | -0.08 | D |
| 77 | A | 0 | 77 | 18 | 0.00 | 0.00 | 2.06 | V |
| 78 | A | 0 | 78 | 5 | 0.00 | 0.00 | 4.38 | I |

**Vav-1 (3BJI)**

| pdb | #chain | int_id | res# | aa | DDG (complex) | DDG (complex,obs) | DG (partner) | |
|---|---|---|---|---|---|---|---|---|
| 61 | C | 1 | 429 | 14 | 0.74 | 0.00 | 0.84 | Q |
| 62 | C | 0 | 430 | 4 | -0.15 | 0.00 | 3.14 | E |
| 63 | C | 0 | 431 | 3 | -0.11 | 0.00 | -0.86 | D |
| 64 | C | 1 | 432 | 20 | 2.59 | 0.00 | 0.83 | Y |
| 65 | C | 1 | 433 | 3 | 2.29 | 0.00 | -0.93 | D |
| 66 | C | 1 | 434 | 15 | 2.25 | 0.00 | -0.35 | R |
| 67 | C | 1 | 435 | 10 | 2.55 | 0.00 | 0.70 | L |
| 68 | C | 0 | 436 | 15 | 0.00 | 0.00 | 3.11 | R |
| 70 | C | 1 | 438 | 10 | 1.45 | 0.00 | 0.55 | L |
| 71 | C | 1 | 439 | 16 | 0.63 | 0.00 | 0.02 | S |
| 72 | C | 0 | 440 | 20 | 0.00 | 0.00 | 4.34 | Y |
| 74 | C | 1 | 442 | 14 | 0.32 | 0.00 | -0.67 | D |
| 75 | C | 0 | 443 | 17 | 0.00 | 0.00 | 1.66 | T |
| 76 | C | 0 | 444 | 3 | 0.00 | 0.00 | -0.36 | D |
| 77 | C | 0 | 445 | 18 | 0.00 | 0.00 | 2.11 | V |
| 78 | C | 0 | 446 | 5 | 0.00 | 0.00 | 3.84 | I |

### Materials and Methods

#### Homology model of GEF-H1 in complex with RhoA

The structure of Akap13 in complex with RhoA (PDB 4D0N) was used as a template to build a GEF-H1 model in complex with RhoA, as the most related GEF with X-Ray data. Due to the localized binding site analysis required in this project, a model by single point mutations was generated. Akap13 side-chains within 5Å of P5 were mutated, and side-chains from both sides were energy-minimized using the MOE software (Amber10:EHT forcefield), leaving any atoms outside the binding site of P5 constrained. Following mutations were introduced (amino acids in the vicinity of P5 are highlighted, with the red ones corresponding to the ones that required mutation in Akap13):

```
FCSRHSKALKLYKELYARDKRFQQFIRKVTRPAVLKRHGVQECILLVTQRITKYPLLISRILQHSHGIEEERQDLTTALGLVKELLSNVDEGIYQLEKGAR
FCGQHNQSVNYFKDLYAKDKRFQAFVKKKMSSSVVRRLGIPECILLVTQRITKYPVLFQRILQCTKDNEVEQEDLAQSLSLVKDVIGAVDSKVASYEKKVR
341                                                                                                441
```

Modeling of stP5 was then performed by energy minimization in rigid GEF-H1.

#### Synthesis of stP5

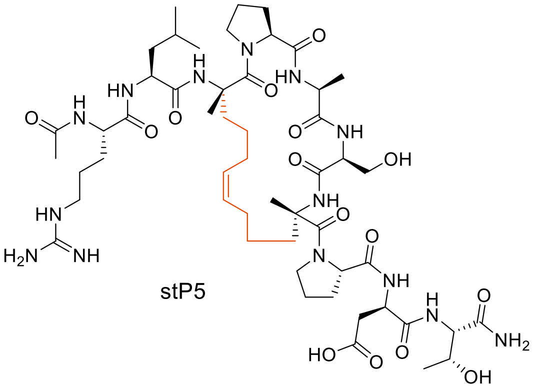

stP5

The following peptide sequences were ordered from Peptide Synthetics on Rink amide resin: Ac-RLXPASXPDT-$NH_2$ and Ac-RLXPLSXPDT-$NH_2$. X=(S)-2-(2'-pentenyl)Ala.

A dried microwave vial was charged with peptide resin (35 w-%, 120 mg, 36 µmol, 1.00 eq) and Hoveyda-Grubbs II catalyst (2.2 mg, 3.6 µmol, 10 mol-%) under an $N_2$ atmosphere. Degassed DCE (1.4 mL) was added, followed by a degassed solution of LiCl in DMF (0.4 M, 0.15 mL). The suspension was stirred and heated to 90°C under microwave irradiation for 1h. The resin was then washed with DMF (3x), $CH_2Cl_2$ (3x), and MeOH (3x) using an SPE column equipped with a frit. The resin was dried through a flow of $N_2$ and was resuspended in TFA/TIPS/$H_2O$ 95/2.5/2.5 (0.5 mL). The suspension was gently shaken for 2h15 in the SPE column, and the resin was then washed with TFA (3x). The filtrate was concentrated using a constant stream of $N_2$ to yield a brown oil. The peptide was precipitated using cold MTBE, and the mixture was centrifuged to collect the precipitate. LC-MS analysis revealed that the main side products are a demethylated specie, a *tert*-butylated specie, and a TFA ester. The latter was easily removed by a minute treatment with 0.15 eq of aqueous 0.1 M LiOH before purification. Purification by flash chromatography (C18, 30-50% MeCN/H2O, 0.1% TFA) and subsequent freeze-drying afforded stP5 as an off-white fluffy solid (8.5 mg, 21% yield).

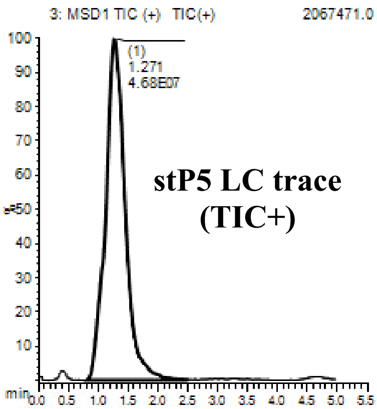

HR-MS (ESI): stP5 $[C_{55}H_{92}N_{14}O_{15}+H]^+$ ($MH^+$) 1147.64698; found 1147.6449 (error -1.82 ppm)

The TAT-stP5 peptide sequence Ac-RLXPLSXPDT-YGRKKRRQRRR-$NH_2$ was further obtained from Eurogentec (95% pure by HPLC) to aid cellular penetration.

#### Circular Dichroism Spectra

CD spectra were collected on a Chirascan Series Spectrometer at 4°C in a 1 mm quartz cuvette. The instrument was calibrated with trifluoroethanol. Each plotted line represents the mean of 10 repeated scans with a 1 nm bandwidth and a time-per-point of 1 s for a nominal step size of 1.0 nm. Data were collected between 180 and 260 nm with appropriate background subtraction.

#### Purification of His-tagged GEF-H1

Amino acids 201-601 of human GEF-H1 DNA were amplified by PCR using the following primers

F: 5' AGGATCGATGGGGATCCGAGCTGATGAGTG`ACTTTGAGATG 3'
B: 5' GCCGGATCAAGCTTCGAATTTTAGTCCTTCTGCTGCAACT 3'

The fragment was gel-purified and cloned into the pRSET-A vector backbone (Invitrogen) using In-Fusion HD cloning (Clontech) at the EcoRI and BamH1 sites in the vector. Plasmids were verified by sequencing.

The plasmid was transformed into chemically competent BL21(DE3) pLysS *E. Coli* cells (NEB C2527H). Starter cultures were grown in Luria-Bertani (LB) broth with ampicillin overnight at 37°C (280 rpm). The next day, cultures were expanded (5 mL/L LB + ampicilin) and grown at

37°C (200 rpm) until $OD_{600}$ = 0.7 was reached. Protein expression was then induced with 1 mM IPTG for 16h at 16°C (200 rpm). Cultures were spun down at 4°C for 20 min (4000 rpm, JS4.5 rotor), and the pellets were resuspended in cold lysis buffer (10 mL/g pellet). The suspensions were then frozen at -80°C until processing. Cell suspensions were defrosted, supplemented with 1 mM PMSF, and sonicated on ice for 10 min in 10s intervals with 20s breaks (Vibra-cell, 60% of maximum intensity). The lysates were clarified at 30 000xg for 30 min at 4°C and filtered (0.45 µm) before loading on a 5 mL HisTrap Ni NTA column (Cytiva, 5 mL/mL) equilibrated with lysis buffer containing 10 mM imidazole. Columns were washed lysis buffer containing 20 mM imidazole, then eluted with a 20 mM-500mM imidazole gradient. Fractions containing the main protein peak were pulled and centrifuged at 17000xg. The supernatant was desalted into a storage buffer using desalting columns (GE PD10). The eluates were concentrated down to 2 mL (Amicon, 10 kDa, 10 mg/mL) and spun down at 17000xg before loading on a size exclusion chromatography column (HiLoad Superdex 75 pg, Cytiva) pre-equilibrated in storage buffer. Eluted fractions were analyzed by SDS-page, pulled, and concentrated to 3.5 mg/mL using concentrating columns (Amicon, 10 kDa) columns. Approximately 6 mg protein per L culture was obtained. The protein was analyzed by SDS-page, immunoblot, MS, and TSA.

Lysis buffer: 50 mM Tris-HCl pH 8.0, 150 mM NaCl, 0.1% Tx100, 10% glycerol, 0.5 mM DTT, lysozyme. Storage buffer: 50 mM Hepes pH 7.5, 125 mM NaCl, 1 mM DTT.

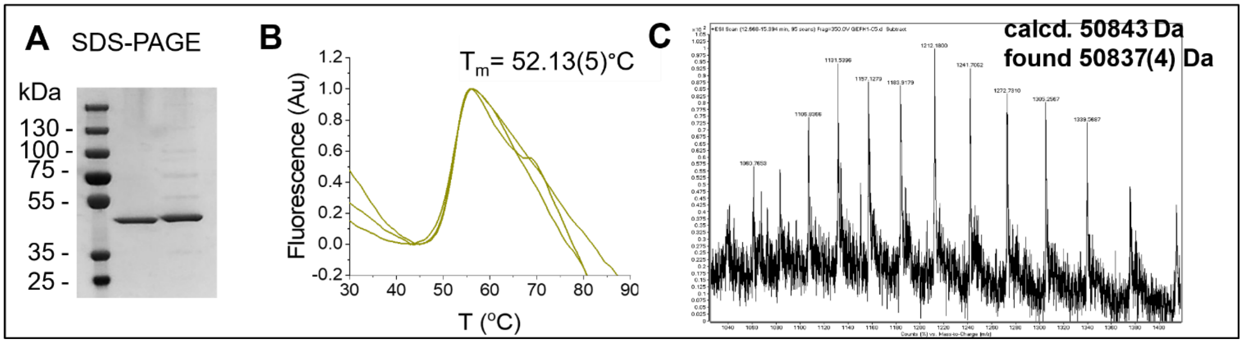

Figure SS1 – Quality control for GEF-H1 fusion protein. **A.** SDS-Page of two different batches. **B.** Melting point determination by TSA in a phosphate buffer (3 duplicates) **C.** MS spectrum (C18 phase LC separation, ESI ionization)

#### Surface plasmon resonance

Compounds were evaluated for their affinity towards GEF-H1 by SPR using a Biacore T200 system at 25˚C. The protein was diluted to 40 µg/mL in 10 mM sodium acetate at pH 5.5-6. Data was collected on a dual flow cell on CM5 amine coupling sensor chips (Cytiva BR100530). With PBSP+ as a running buffer, the surface was initially activated with 1:1 N-hydroxysuccinimide (0.1 M) and 1-Ethyl-3-(3-dimethylaminopropyl)carbodiimide (0.4 M) for 420s at 10 µL/min. Protein solutions were pulsed on the surface through the automatic immobilization tool (Biacore T200 software), and any unreacted surface was quenched by 420s of ethanolamine (1M, pH 8.0). The blank reference channel was prepared in the same way omitting the injection of protein. Chips with immobilization between 2000 RU and 8000 RU were prepared depending on the molecular weight of the analyte and immobilization efficiency. Equilibrium responses against concentration were plotted and analyzed by non-linear regression (Origin software fit equation RI + Rmax * x / (KD + x)) to obtain binding affinities ($K_D$). P5: A 4200 RU chip was pre-equilibrated using HBS-P+ as a running buffer. Serial dilutions of P5 in HBS-P+ were injected in duplicates over the surface for 60 s and left to dissociate for 200 s at a 30 µL/min flow rate. stP5:A 8000 RU chip was pre-equilibrated using PBS-P+ 2% DMSO as a running buffer. Serial dilutions of stP5 in PBS-P+, 2% DMSO, were injected in duplicates over the surface for 60 s and left to dissociate for 100 s at a 30 µL/min flow rate. A 50% DMSO wash of the tubing was added at the end of each injection.

#### Cell culture

MDCK cells were cultured in high glucose Dulbecco's Modified Eagle Medium (DMEM) containing 10% fetal bovine serum (FBS) and 1% Penicillin/Streptomycin. The cells stably transfected with GEF-H1-VSV, under the control of a doxycycline-inducible manner were described previously[2, 3]. The MDCK cells were maintained in the medium supplemented with 5 µg/mL Blasticidin (PAA Laboratories) and 400 µg/mL Zeocin (Invitrogen, Waltham, MA, USA). Doxycycline was added at 4 µg/mL to induce construct expression. Primary HDMEC (PromoCell) cells were cultured in endothelial cell growth medium MV2 (PromoCell, Heidelberg, Germany) supplemented with C-39225 supplement mix (PromoCell) and cells were used between passage two and four as described previously[4, 5].

#### Immunocytochemistry

Cells were fixed with 3% PFA in PBS for 15 min at room temperature followed by permeabilization with 0.3% Triton X-100 in PBS for 10 min at room temperature. Cells were then blocked for 15 min with blocking buffer (1% BSA, 0.1% NaN3, 20 mM glycine, PBS). Cells were stained with the following primary antibodies overnight at 4°C in blocking buffer: mouse anti-ZO-1 (Invitrogen 33-9100) followed by secondary antibody incubation for 1 h at room temperature using FITC Donkey anti-mouse (Jackson Immunoresearch) plus Hoechst. Cells were mounted on slides using Prolong Gold Antifade. Samples were imaged using a Nikon Live cell microscope.

#### Gene reporter assays

MDCK with inducible expression of VSV-GEF-H1 cells were seeded in 96-well plates the day before transfection at 5000 cells per well. Cells were transfected with a plasmid containing an NF-kB promoter driving firefly luciferase expression and a plasmid containing a reference CMV promoter driving renilla expression using TransIT-X2 transfection reagent (Mirus, MIR 6000). Cells were then incubated with Doxycycline (2 ug/ml) in the absence or presence of GEF-H1 inhibitors overnight. Quadruplicate wells were used for each condition in all experiments. The next day, luminescence values were measured using a dual luciferase assay kit (Promega Corp, Madison, WI, USA) and a FLUOstar OPTIMA microplate reader (BMG Labtech, Ortenberg, Germany).

#### Cell shape data analysis

Cell images of immunofluorescent staining (ZO-1 and Hoechst) were visualized in Napari. The cells were segmented using Serialcellpose, a Napari plugin (https://www.napari-hub.org/plugins/napari-serialcellpose) to batch segment cells with Cellpose 2 using custom models that we trained for MDCK and HDMEC cell segmentation[6]. Napari and cellpose were installed as Python packages and managed using Anaconda. After cell segmentation, the segmented cell images were analyzed by Fiji (ImageJ) and AR ratios of cells were calculated. AR ratio of a cell was defined as a ratio of cell length (L) to cell width (W) which could indicate cell shape changes such as cell elongation. Untreated AR values were deducted from experimental AR values and then divided by the corresponding value obtained from Doxycline-treated cells. The resulting value was then multiplied by the AR standard deviation normalized to the untreated sample standard deviation. The resulting value was named AR index.
